# FIRST INSIGHTS IN A NON-RODENT MODEL SPECIES OF THE OLFACTORY LIMBUS. THE RED FOX (*Vulpes vulpes*) AS A CASE IN POINT

**DOI:** 10.1101/2022.11.08.515585

**Authors:** Irene Ortiz-Leal, Mateo V. Torres, Víctor Vargas-Barroso, Luis Eusebio Fidalgo, Ana López-Beceiro, Jorge Larriva-Sahd, Pablo Sanchez-Quinteiro

## Abstract

The mammalian olfactory systems can be divided into several subsystems based on the anatomical location of their neuroreceptor cells and the family of receptors they express. The more in depth studied systems are the main olfactory system and the vomeronasal system, whose first integrative enters are the main and the accessory olfactory bulb, respectively. In addition, there is a range of olfactory subsystems which converge to the transition zone located between the main olfactory bulb and the accessory olfactory bulb., which has been termed as olfactory limbus (OL) and includes specialized glomeruli which receive uncanonical sensory afferences and interact with the MOB and AOB. Beyond the laboratory rodents, there is a lack of information regarding the olfactory subsystems of carnivores. We have focused on the specific study of the olfactory limbus of the fox, performing serial histological sections, general and specific histological stainings, including both double and simple immunohistochemical and lectin-histochemical labeling techniques. As a result, we have been able to determine that the OL of the fox shows an uncommon development with a high degree of development and complexity. This makes this species a novel mammalian model that could provide a wider understanding of non-canonical pathways involved in the processing of chemosensory cues.

## INTRODUCTION

Nowadays, neuroanatomical, electrophysiological, and genomic studies have shown that the concept of a single olfactory system is an oversimplification. The seemingly simple mammalian nasal cavity hides within it a significant number of olfactory systems, some of them only relatively recently discovered (Barrios et al., 2014a). The mammalian olfactory systems can be divided into several subsystems based on the anatomical location of their neuroreceptor cells, the family of receptors they express, the signaling mechanisms employed in their transduction chain, the chemosensory stimuli they detect, and the axonal targets of their sensory neurons in the rhinencephalon (Munger, 2009). However, a growing number of evidence supports a synergic and cooperative interaction between all of them (Mucignat-Caretta et al., 2012; Pardo-Bellver et al., 2022).

The more in depth studied systems are the main olfactory system (MOS) and the vomeronasal system (VNS), both present, with few exceptions, in most mammalian groups. In the MOS the olfactory receptors (OR) are located in the cilia of the mucosa lining the ethmoidal turbinates and in the caudal part of the nasal septum (Salazar et al., 2019). The axons of the neuroepithelial cells project to the glomeruli of the main olfactory bulb (MOB) (Crespo et al., 2013). The MOS responds to thousands of volatile chemosignals which carry information regarding food, pathogens, prey, predators, or conspecifics (Firestein, 2001).

VNS receptors are located in the microvilli lining the neuroepithelium of the vomeronasal organ (VNO) (Salazar et al., 1997, 2003). The vomeronasal neurons send their axons to the accessory olfactory bulb (AOB) where those mammalian groups expressing the two families of vomeronasal receptors V1R and V2R establish a morphofunctional antero-posterior subdivision (Torres et al., 2020, 2022). The sensory neurons of the VNO detect a range of non-volatile natural ligands present in the exocrine secretions of conspecifics, which are involved in innate socio-sexual behaviors (Pardo-Bellver et al., 2017; Pallé et al., 2020; Villafranca-Faus et al., 2021).

In addition, a range of sensory olfactory subsystems have been characterized, mainly in laboratory rodents, such as the Grüneberg ganglion, the septal organ, and the guanylyl cyclase-D-expressing chemosensory neurons (Zimmerman and Munger, 2021). The projection areas of the chemosensory neurons that form these systems configure several anatomical domains in the olfactory bulb. The Grüneberg ganglion (GG) was first described by Hans Grüneberg (1973), and consists of olfactory-marker-protein (OMP) immunopositive cells located at the dorsal area of the nasal vestibule. These cells do not possess prominent dendrites, cilia, or microvilli and lack direct access to the vestibular surface (Breer et al., 2006). Therefore, GG cells could be involved in the detection of highly membrane permeant stimuli. GG cells project to the dorsal regions of the caudal MOB, near the AOB (Fuss et al., 2005; Storan and Key, 2006). The septal organ of Masera (SO) is an isolated patch of sensory epithelium present in Rodentia and located near the base of the nasal septum at the entrances to the nasopalatine ducts (Adams, 1972). The SO expresses 50–80 genes of the OR family, all of them expressed in the MOE as well (Kaluza et al., 2004). OSNs in the SO project to a small cluster of glomeruli located in the caudal, ventro-medial aspect of the MOB (Levai et al., 2003).

The guanylyl cyclase-D-expressing chemosensory neurons (GC-D+) are a subpopulation of olfactory neurons that project to a well-defined domain of the olfactory bulb: the most caudal glomeruli of the MOB, named necklace glomeruli. The term necklace glomeruli was adopted by Shinoda et al. (1993), who identified in the rat a subset of OSNs immunoreactive to human placental antigen X-P2 (PAX) and which converged on 7–9 glomeruli aligned along the caudal MOB. These glomeruli overlapped with a subset of “atypical” glomeruli that displayed acetylcholinesterase (AChE) reactivity (Zheng et al., 1987; Weruaga et al., 2001). The aforementioned projections of the septal organ and Grüneberg’s ganglion overlap as well with the necklace complex. This receptor system remains minimally studied and poorly understood (Zimmerman and Munger, 2021), but it is known that GC-D+ neurons surprisingly are not OMP+, and function as receptors for uroguanylin, guanylin, and components of urine (Leinders-Zufall et al., 2007). The *Gucy2d* ortholog is in most primates a pseudogene but it is intact in some prosimians and in dog, mouse, rat, and tree shrew. Thus, GC-D+ neurons may maintain a chemosensory role in these mammals (Young et al., 2007). Finally, it is also worth mentioning the MS4A family of chemosensors, which are also expressed in the necklace glomeruli, are non-GPCRS, unlike most other types of sensory neurons, several different receptors can be expressed in the same cell, and have been shown to be behaviourally relevant (Greer et al., 2016).

All these olfactory subsystems converge to the transition zone located between the main olfactory bulb and the accessory olfactory bulb. This area consists of a modified bulbar cortex bounded anteriorly by the dorsal MOB and posteriorly by the anterior AOB (Larriva-Sahd 2012). This area has been termed olfactory limbus (OL), and includes specialized glomeruli which receive uncanonical sensory afferences and interact with the MOB and AOB, opening the possibility that OL is a site of non-olfactory and atypical vomeronasal sensory integration (Vargas-Barroso et al., 2017).

There is a lack of information regarding the olfactory subsystems of carnivores. No sensory systems equivalent to the septal organ, Grüneberg’s ganglion and necklace glomeruli have been described in either dogs or cats (Salazar and Sánchez-Quinteiro, 2011; Barrios et al., 2014b). However, studies that have addressed the characterization of the accessory olfactory bulb in mink, meerkats, and dog have pointed to a more complex glomerular organization in the OB than described in other mammalian orders. Thus, the presumptive AOB of mink comprises not only a main dorsocaudal protuberance but also smaller lateral and medial regions (Salazar et al., 1998), whereas a recent study of the meerkat OB found in the vicinity of the AOB a subpopulation of atypical glomeruli with strongly CR-positive neuropil (Torres et al., 2021). Finally, the study of Miodonski in the dog (1968) considered that the AOB consisted of distinct glomerular aggregations; both “in the dorsomedial side of the olfactory bulb, at the posterior margin of the main olfactory bulb and downward along the caudal edge of the main olfactory bulb, descending to its base both on the medial and on the lateral side.” Miodonski’s observations could be interpreted as an erroneous attribution to the accessory olfactory system of atypical glomerular structures belonging to a different olfactory subsystem. However, this multiform characterization of the dog AOB was not confirmed by subsequent lectin and immunohistochemical studies (Nakajima et al., 1998; Salazar et al., 2013), except for the fact that Nakajima et al. (1998) found a small group of glomeruli with NADPH-diaphorase reactivity “between the glomeruli at the most caudal portion of the MOB”. They attributed these glomeruli to the specific projection from subsets of neurochemically different olfactory receptor cells.

Recent studies of the AOB of wild canids performed in the African wild dog (Chengetanai et al., 2020) and especially in the red fox (Ortiz-Leal et al., 2022b) have shown the existence of significant differences between the AOB of domestic and wild canids. Furthermore, the latter study described the presence of a small atypical glomerular formation in the proximity of the AOB. For this reason, in this work we have focused on the specific study of the olfactory limbus of the fox, performing serial histological sections, general and specific histological stainings, including both double and simple immunohistochemical and lectin-histochemical labeling techniques. As a result, we have been able to determine that the OL of the fox shows an uncommon development with a high degree of development and complexity. This makes this species a novel mammalian model that could provide a wider understanding of non-canonical pathways involved in the processing of chemosensory cues.

## MATERIAL AND METHODS

### 1.1. Samples

For this investigation, 3 male and 2 female adult foxes were employed. They were obtained through hunting expeditions led by the Galician Hunting Federation with the required authorizations granted by the Galician Environment, Territory and Tenement Council. The animals were brought to the Veterinary Faculty of Lugo’s facilities as soon as they were shot, in the field, with no more than a two-hour interval. There, the rostral section of the encephalon was removed using an electric plaster cutter and a gouge clamp and preserved in Bouin’s fixative (Bf). Afterwards, the olfactory bulbs in conjunction with the rostral frontal lobes were embedded in paraffin wax and serially cut by a microtome in a horizontal plane along its entire length with a thickness of 6-7 µm. Hematoxylin-eosin and Nissl stains, lectin histochemistry, and both single and double immunohistochemistry were used to stain the slides.

### 1.2. Lectin histochemistry staining

*Ulex europaeus* agglutinin lectin (UEA) was employed as a first step in some of the sequential double immunohistochemical labelings. UEA labels the VNS pathway in several species, including the fox (Ortiz-Leal et al., 2022b) and dog (Salazar et al., 1994).

All sample slides were deparaffinized and rehydrated prior to beginning the lectin histochemistry procedure. The samples were then incubated to 3% H_2_O_2_ solution for 30 minutes to inhibit endogenous peroxidase activity, followed by two rinses in pH.7.2, 0.1M phosphate buffer (PB), which blocked non-specific binding. The slides were then washed three times for five minutes in a PB solution, followed by an incubation period of 1 hour at room temperature in a 0.5% BSA/UEA solution. A further overnight incubation with an anti-UEA peroxidase-conjugated antibody was performed on the slides. The samples were rinsed in 0.2 M Tris-HCl, pH 7.61 for 10 minutes the next day, followed by a PB wash, before being developed using a DAB chromogen. Controls involved removing the UEA and allowing the lectin to be preabsorbed by employing an excessive quantity of the matching sugar, L-fucose.

### 1.3. Simple immunohistochemical staining

The fox olfactory limbus was examined in-depth using immunohistochemistry. Among the antibodies employed (Table 1), the anti-Gαo and anti-Gαi2 antibodies are particularly useful because they label the transduction cascade for V2R and V1R vomeronasal receptors, respectively. Using antibodies against microtubule-associated protein 2 (MAP-2), neuronal dendritic development in the olfactory limbus was studied. Prior research has shown that the distribution of calcium-binding proteins may be used as a neuronal marker to distinguish between distinct brain areas and neuronal subpopulations (Baimbridge et al., 1992). Therefore, the calcium-binding proteins calbindin (CB), calretinin (CR), and secretagogin (SG), which play a role in controlling the levels of cytosolic free calcium ions in neurons, were studied using immunohistochemistry.

**Table 1.**
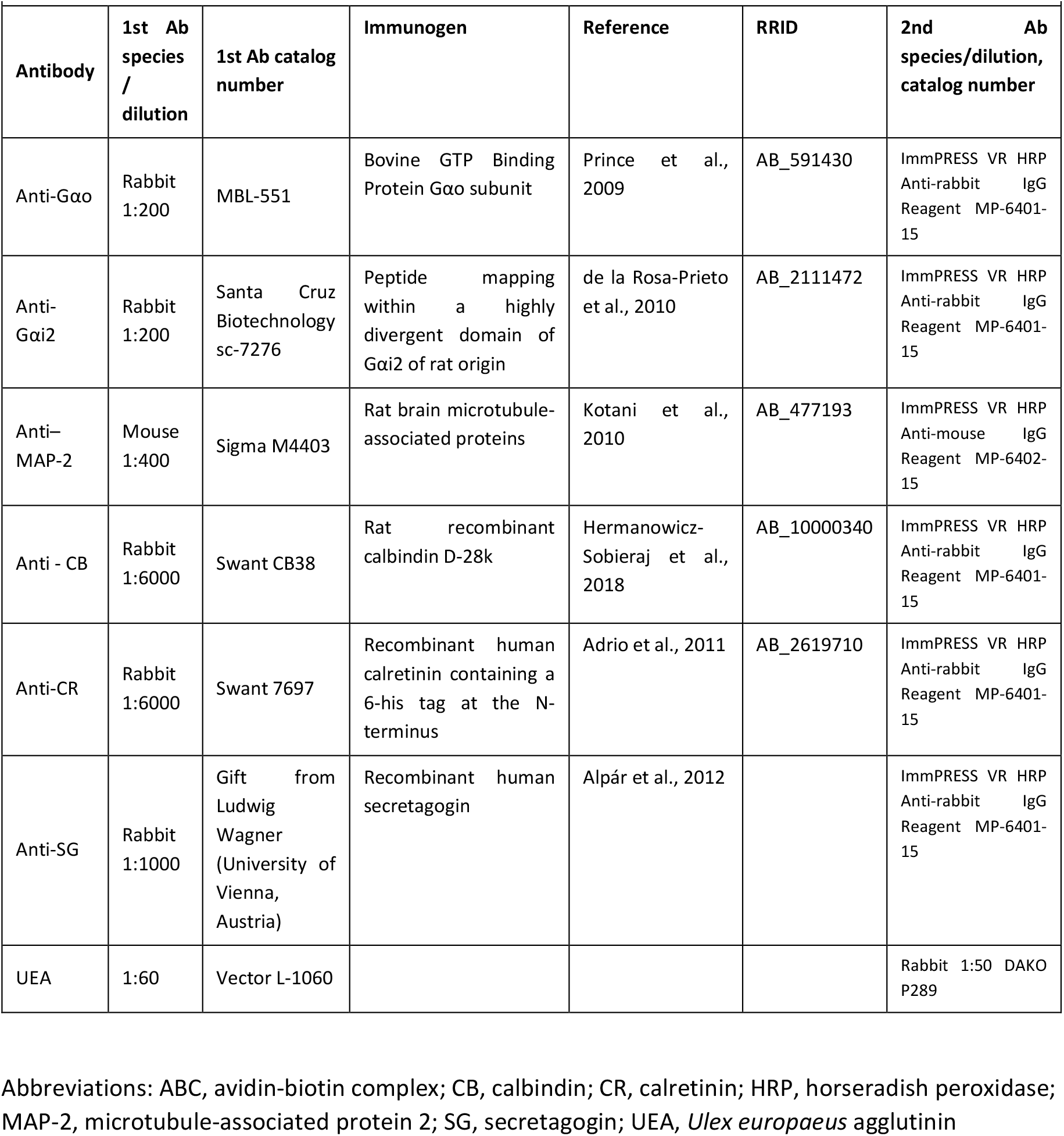
Detailed information on the antibodies and lectins used in this study.

#### 1.3.1. Antibody characterization and specificity

Table 1 provides details for all antibodies, including their suppliers, dilutions, target immunogens, and Research Resource Identifiers (RRID). In every instance, the immunostaining patterns produced in the red fox using these antibodies matched those previously obtained in a number of mammalian species. Table 1 lists relevant references for each antibody.

#### 1.3.2. Simple immunohistochemical protocol

Deparaffinized and rehydrated samples were treated for 15 min in a 3% H_2_O_2_ solution to inactivate endogenous peroxidase activity prior to the immunohistochemistry reaction. To prevent non-specific binding sites, 2.5% horse normal serum from the ImmPRESS reagent kit Anti-mouse IgG/Anti-rabbit IgG (Vector Laboratories, Burlingame, CA, USA) was used for 30 min (Table 1). After that, the samples were treated with the primary antibody for an overnight incubation at 4°C with humidity. Using either the ImmPRESS VR Polymer HRP Anti-Rabbit IgG or the Anti-Mouse IgG reagent, samples were incubated for 30 min (Table 1). In every instance, 3 × 3 minute PB washes were carried out in between steps. Finally, slides were developed with a DAB chromogen (using the same procedure as for the lectin histochemical labeling) and then dehydrated and mounted.

The omission of the primary antibody was employed as a negative control for all immunohistochemistry methods, and none of the negative control samples showed any labeling or non-specific background staining. We repeated the immunohistochemistry process on previously unstained mouse or rabbit tissue from earlier investigations as a positive control. These samples were all known to express the desired proteins, and each time, the expected positive results were obtained.

#### 1.3.3. Double-immunohistochemical protocol

A sequential twice-repeated enzyme-labeled approach was used for double immunostaining (Hasui et al., 2003). The sections were given a 5-minute dip in 0.1 M glycine solution (pH 2.2) in between both immunolabelings. To select the most suitable dye to visualize the immunoreaction, both DAB and Vector VIP Peroxidase Substrate Kit (SK-4600, Vector Laboratories) were combined exchanging their order. Using first DAB and then VIP was the optimal combination for our immunostaining.

### 1.4. Image acquisition and digital processing

A Zeiss Axiophot microscope and a Karl Zeiss Axiocam MRc5 digital camera were used for image acquisition. Using Adobe Photoshop CS4 the white balance settings were adjusted (Adobe Systems, San Jose, CA, USA). No particular details were added, relocated, improved, or removed from the photographs.

## RESULTS

The microscopic study of the olfactory limbus (OL) of the fox was performed on serial horizontal sections of the anterior region of the telencephalon, across an area that included the accessory (AOB) and main olfactory bulbs (MOB) and the frontal lobe of the brain (FL). The term olfactory limbus refers to the transition zone between the MOB and the AOB; a large area located in the medial part of the olfactory bulb, along the area delimited between the dorsocaudal edge of the MOB and the anterior end of the AOB (Fig. 1). It is not a sharp frontier, but it is rather a heterotypical bulbar cortex, whose cytological characteristics differ from those of the MOB and AOB. The most obvious characteristic of the OL is the variable extent of modification from the laminar pattern observed throughout the olfactory bulb. The widespread presence of atypical glomeruli in the olfactory limbus of the fox species is remarkable (Figs. 1B, C). In contrast to the linear arrangement and uniformity of size of the MOB glomeruli (100-150 µm maximum diameter), OL glomeruli are characterized by an atypical, disarrayed arrangement, an irregular shape and a wide diversity of sizes (50-500 µm maximum diameter).

**Figure 1.**
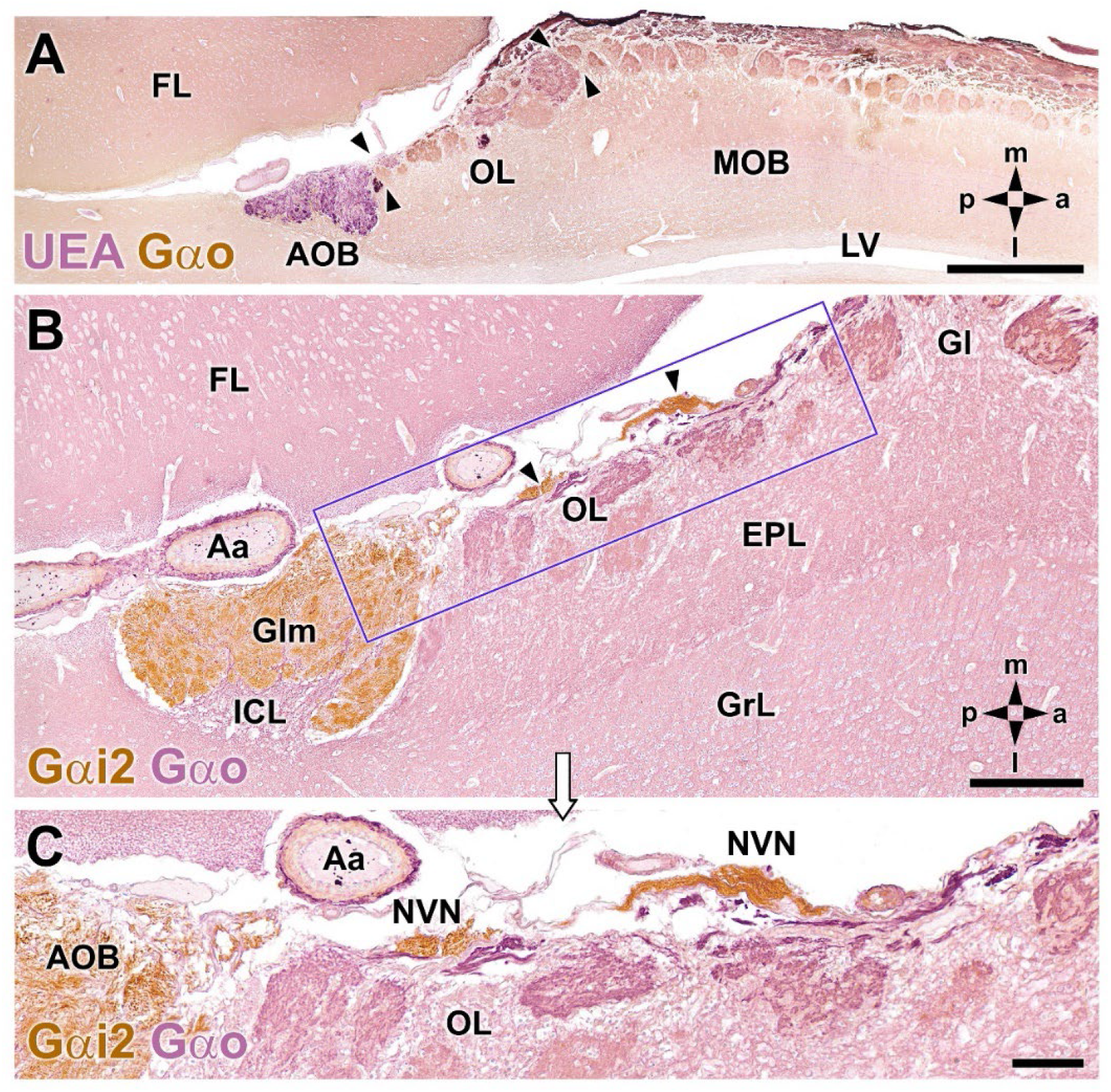
Double-immunohistochemical labeling of the fox olfactory limbus (OL). **A**. Double immunostaining with the lectin UEA (magenta) and Gαo antibody (brown). The vomeronasal nerve and glomerular layers of the AOB are strongly labeled with UEA. Anti-Gαo immunolabeling shows a widespread immunopositive pattern, more intensely in both the olfactory nerve and glomerular layers of the MOB. The area of the olfactory limbus, delimited by arrowheads comprises irregularly shaped glomeruli, without a homogeneous pattern of immunostaining. **B** and **C**. Double immunostaining for Gαi2 (brown) and Gαo (magenta). Anti-Gαi2 stains the superficial AOB and the *nervus vomeronasalis* (NVN) (arrowheads). Anti-Gαo stains the internal cellular layer (ICL) of the AOB and the neuropil of the OB. The olfactory limbus, box enlarged in **C**, comprises irregularly shaped glomeruli without a homogeneous pattern of immunostaining. The branches of the NVN, Gαi2 immunopositive, contrast with the nerve endings reaching the glomeruli of the olfactory limbus which are Gαo immunopositive. Aa, Artery; EPL, external plexiform layer; FL, frontal lobe; Gl, MOB glomerular layer; Glm, AOB glomerular layer; GrL, MOB granular cell layer; LV, lateral ventricle. The compass indicates the orientation, as follows: m, medial; l, lateral; p, posterior; a, anterior. Scale bars: A: 1cm. B, C: 250 µm.

Double immunohistochemical labeling against different markers allowed us to characterize the topographical relationships between the atypical glomeruli (Fig. 1A), the AOB and the axonal endings of the vomeronasal nerve (NVN) (Fig. 1B). Double immunolabeling against the UEA lectin and the G-protein subunit Gao (Fig. 1A) allows the identification of the two superficial layers of the AOB, the vomeronasal nerve and glomerular layers, both of them intensely stained with the lectin. Conversely, anti-Gao staining was observed in the deep layers of the AOB and in the whole MOB, although more intensely in both the olfactory nerve and glomerular layers. It is striking how the area of the olfactory limbus, delimited by arrowheads in Fig. 1A, comprises irregularly shaped glomeruli, without a homogeneous pattern of immunostaining.

Double immunolabeling against the G protein subunits, Gao and Gai2, (Fig. 1B) is equally useful for the study of this area, as Gai2 expression is exclusive to the vomeronasal system, both at the level of the surface layer of the AOB and the NVN (Fig. 1C). In contrast to the NVN, the nerve endings reaching the glomeruli of the olfactory limbus are immunopositive to Gao (Fig. 1C).

The histological study with both hematoxylin-eosin (Fig. 2A) and Nissl staining (Fig. 2B) allowed to recognize the texture of the olfactory limbus. It is characterized by a high structural complexity. The two most striking discernible features are the presence of dense and compact clusters of neuronal somata (Fig. 2A1, B1) and a broad nervous formations consisting of neuronal somas, not so densely aggregated, distributed within a neuropil which has clearly distinct boundaries from the underlying glomeruli (Fig. 2A2, B2). We have termed this previously undescribed nervous formation as a macroglomerular complex (MGC). In both cases it is remarkable how this high concentration of neuronal somata is located in the most superficial layer of the bulb, in proximity to the pial surface. Deep to both structures, small, irregularly shaped glomeruli complete the organization of the olfactory limbus.

**Figure 2.**
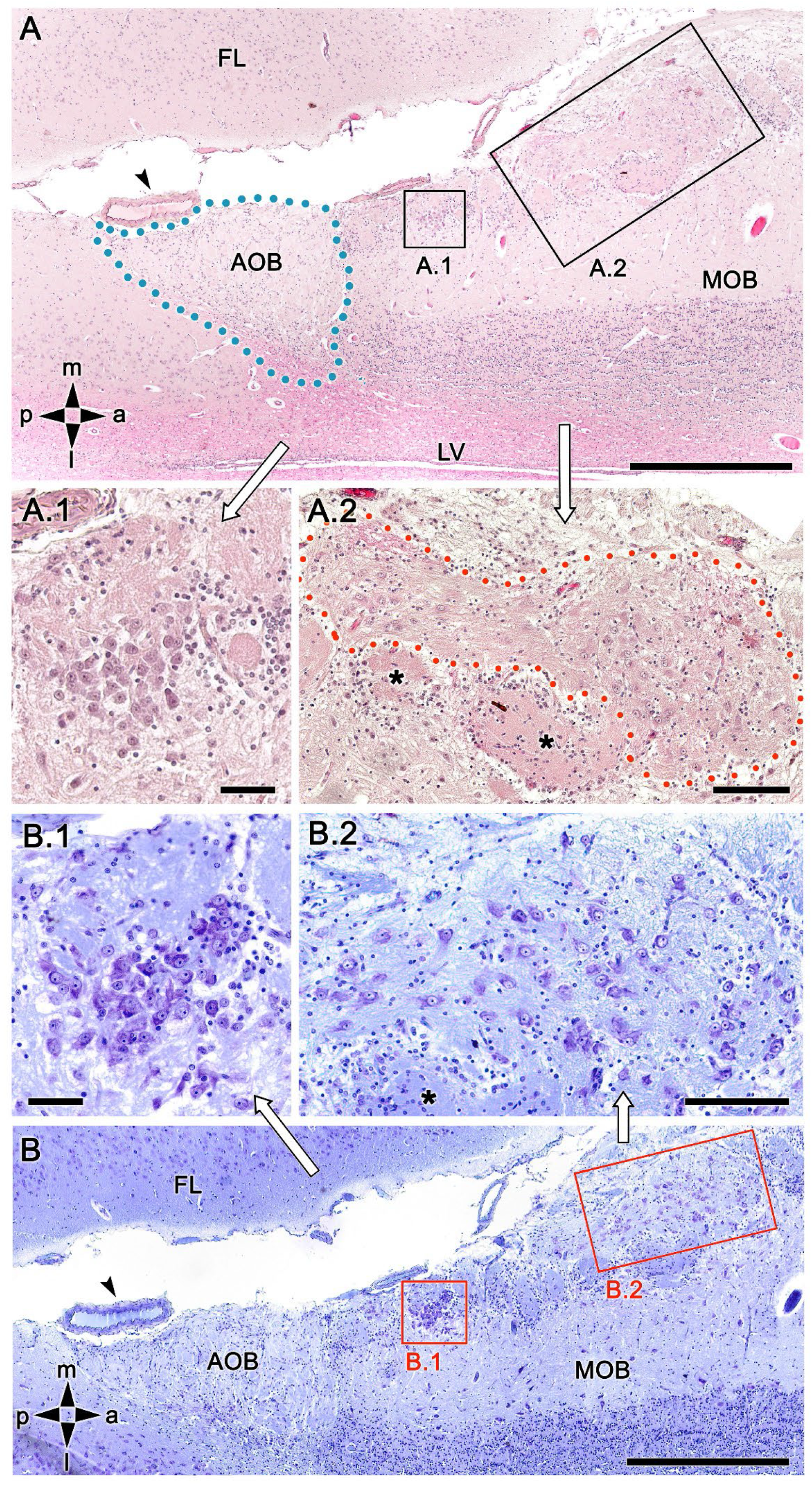
Histological study of the fox olfactory limbus. **A**. General view of the olfactory limbus, cut in the horizontal plane. The accessory olfactory bulb (AOB), close to a large artery (arrowhead) is framed by dots. The two most striking discernible features of the OL are framed and showed at higher magnification in figures **A.1** and **A.2**. They are the dense neuronal cluster (A.1) and the macroglomerular complex: a broad nervous formation consisting of neuronal somata distributed within a neuropil with clearly distinct boundaries (A.2) delimited by red dots. Deeper to both structures there are small, irregularly shaped glomeruli (asterisks). **B**. Nissl-stained serial consecutive section showing the morphology of the neuronal somata. In both instances, the denser aggregates (B.1) and the MGC (B.2), possess somata polygonal, ellipsoidal and rounded in shape. FL, frontal lobe; LV, lateral ventricle; MOB, main olfactory bulb. Orientation: m, medial; l, lateral; p, posterior; a, anterior. Scale bars: A, B: 500 µm. A.1, B.1: 100 µm. A.2, B.2: 250 µm.

To rule out that the presence of these atypical formations is the result of interindividual anatomical variability, we extended the histological study with Nissl staining to four more animals. The results of this study are shown in Figs. 3 and 4. In the first of these animals (Fig. 3), the study of the OL in horizontal sections at three levels dorsally to the AOB, allowed us to characterize in more detail the appearance and dimensions of both the neuronal clusters (Fig. 3A, A.1) and the broad macroglomerular formation (Fig. 3A, A.2, B, B.1, C). Somata in the clusters were similar in size, and had either polyhedral, ellipsoid, or oval morphology (Fig. 3A.1, 3A.2, B.1, C). The magnitude of the MGC is noteworthy, as in some sections it extended over several millimeters, always located on the bulbar surface, in close relation to populations of glomeruli located in the immediately deeper plane (Fig. 3A, B). Most of these glomeruli had irregular, atypical shapes (Fig. 3A, B), although sometimes their shape was typically spherical, and their boundaries were well defined (Fig. 3A). Nissl staining of the other three individuals (Fig. 4A, B, C) verified the above findings in all cases.

**Figure 3.**
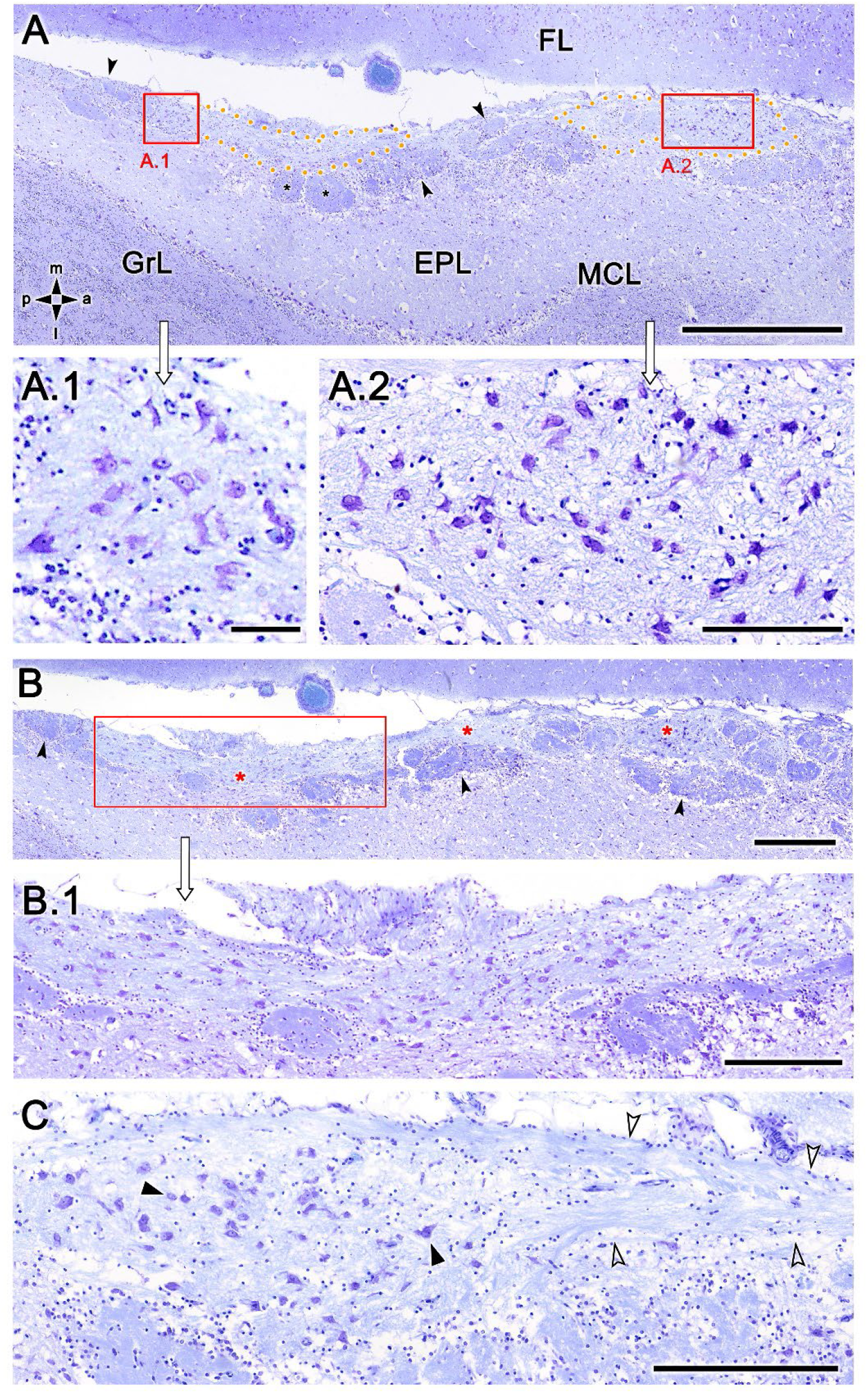
Nissl-stained sections of the fox olfactory limbus at different horizontal levels. The serial study carried out on this individual depicts the development of the olfactory limbus in this species. **A**. Section at a ventral level showing the presence of a neuronal cluster (box enlarged in **A.1**), and the superficial MGC (encircled by yellow dots and partially enlarged in **A.2**.). Most of the glomeruli close to the MGC had an irregular and atypical shape (arrowheads), but some of them were more typically spherical (black asterisk). **B**. Image corresponding to a more dorsal level than the previous one, showing the extent of the MGC (red asterisks), of which a sample (red box) is shown at a higher magnification in **B.1**, and the presence of atypical glomeruli at a deeper level (arrowheads). **C**. A third section at a more dorsal level shows the morphology of the neurons present in the MGC (black arrowheads) and how these neurons are associated with a prominent fascicle of nerve fibers (open arrowheads). FL, frontal lobe of the telencephalon; GrL, granular layer; EPL, external plexiform layer; MCL, mitral cell layer. Orientation: m, medial; l, lateral; p, posterior; a, anterior. Scale bars: A: 1 mm. B: 500 µm. A.2, B.1, C: 250 µm. A.1: 100 µm.

**Figure 4.**
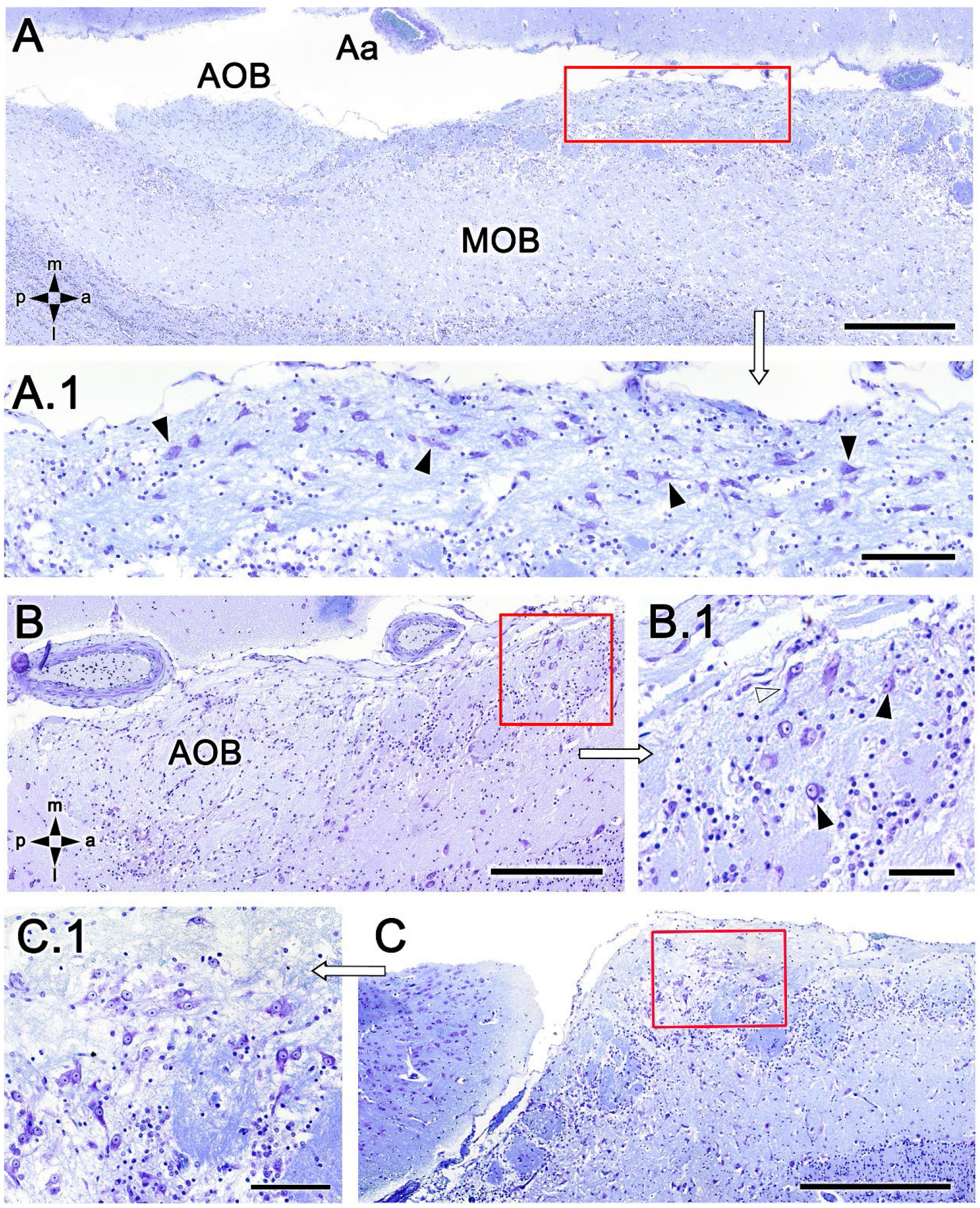
Nissl-stained sections of the fox olfactory limbus from three different individuals. **A**. Section at a ventral level showing the topography of the nervous formation, located on the surface of the bulb, superficial to the glomeruli. **A.1**. At higher magnification (red box in A) numerous neuronal somata are observed (arrowheads). **B**. Horizontal section at the level of the AOB. Anteriorly to the AOB a superficial neuronal cluster, surrounded by atypical glomeruli, is noticeable (red box). **B.1** The neuronal somata have oval shape (arrowheads) and the origin of thick processes is visible (open arrowhead). **C**. In this animal the neuronal cluster is located in a more anterior position, but also very close to the pial surface. **C.1**. At higher magnification the neurons show a similar morphology to that of the previous animal depicted. Aa, artery; AOB, accessory olfactory bulb; MOB, main olfactory bulb. Orientation: m, medial; l, lateral; p, posterior; a, anterior. Scale bars: A, C: 500 µm. B: 250 µm. A.1, C.1: 100 µm. B.1: 50 µm.

The information obtained from the macroscopic study of the olfactory bulb and the interpretation of the histological series is summarized in Figure 5. The macroscopic image of the olfactory bulb shows the area in which the formations described in the olfactory limbus of the fox are located, mainly in the caudomedial border of the MOB (Fig. 5A) and their relationship with the projection area of the AOB. Figs. 5B and C show a schematic drawing of a histological section of the MOB together with an enlargement of the caudodorsal area, showing the irregular organization of atypical glomeruli, both in terms of their arrangement and dimensions, as well as the structure of the macroglomerular complex that runs through it (Fig. 5C).

**Figure 5.**
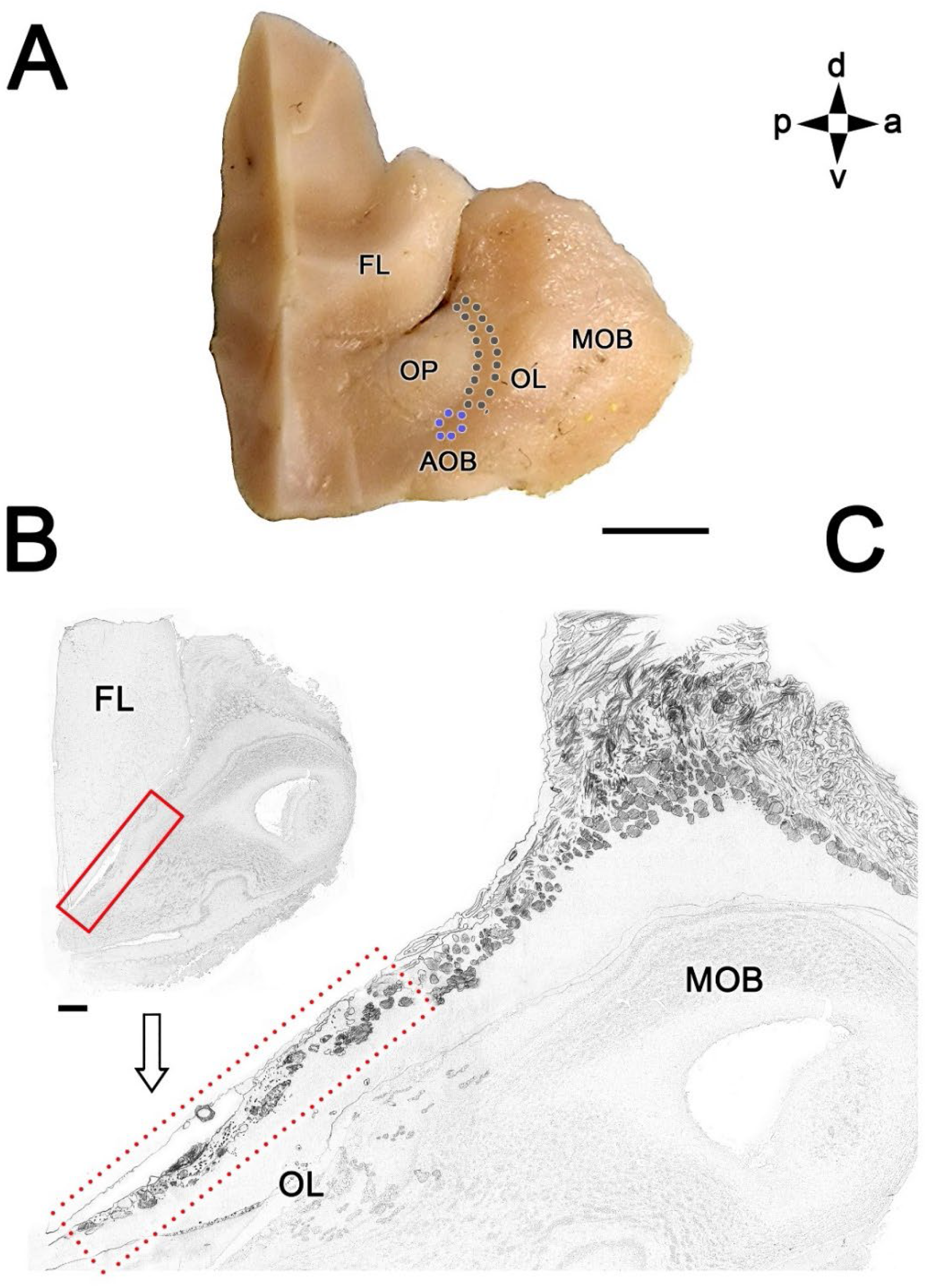
Topographical anatomy of the olfactory limbus of the fox. **A**. Medial view of the left olfactory bulb (MOB), after removal of the medial portion of the frontal lobe (FL). The olfactory peduncle (OP) and the caudo-medial margin of the olfactory bulb are visualized. The accessory olfactory bulb (AOB, blue dots) and the olfactory limbus (OL, black dots) are located along its surface. **B**. Schematic drawing of a sagittal histological section of the olfactory bulb. **C**. The area corresponding to the olfactory limbus (box in B) is enlarged in Figure C. Orientation: m, medial; l, lateral; p, posterior; a, anterior. Scale bars: A: 500 µm. B: 100 µm.

To further characterize the fox olfactory limbus neurochemically and thus determine the possible existence of distinctive features compared to both the accessory and main olfactory bulb, we carried out a specific study with various antibodies and the lectin UEA.

Immunolabeling against the G-protein subunit Gao produced the well-known pattern of immunopositive labeling throughout the neuropil of the frontal lobe of the telencephalon and the olfactory bulb, in which only there is absence of Gao in both the nerve and glomerular layer of the AOB (Fig. 6A). However, it is noticeable that the atypical glomeruli, close to the AOB, are more intensely immunolabeled than the typical glomeruli of the MOB. As for the MGC, it shows a similar intensity of labeling to typical glomeruli. Interestingly, when observing this formation at higher magnification (Fig. 6B), two clearly differentiated areas can be seen in this section; one consisting of numerous neuronal somata and the other devoid of them. These somata are immunopositive for Gao, which contrasts with the somas of the mitral cells of the MOB, which are immunonegative for this marker (Fig. 6C).

**Figure 6.**
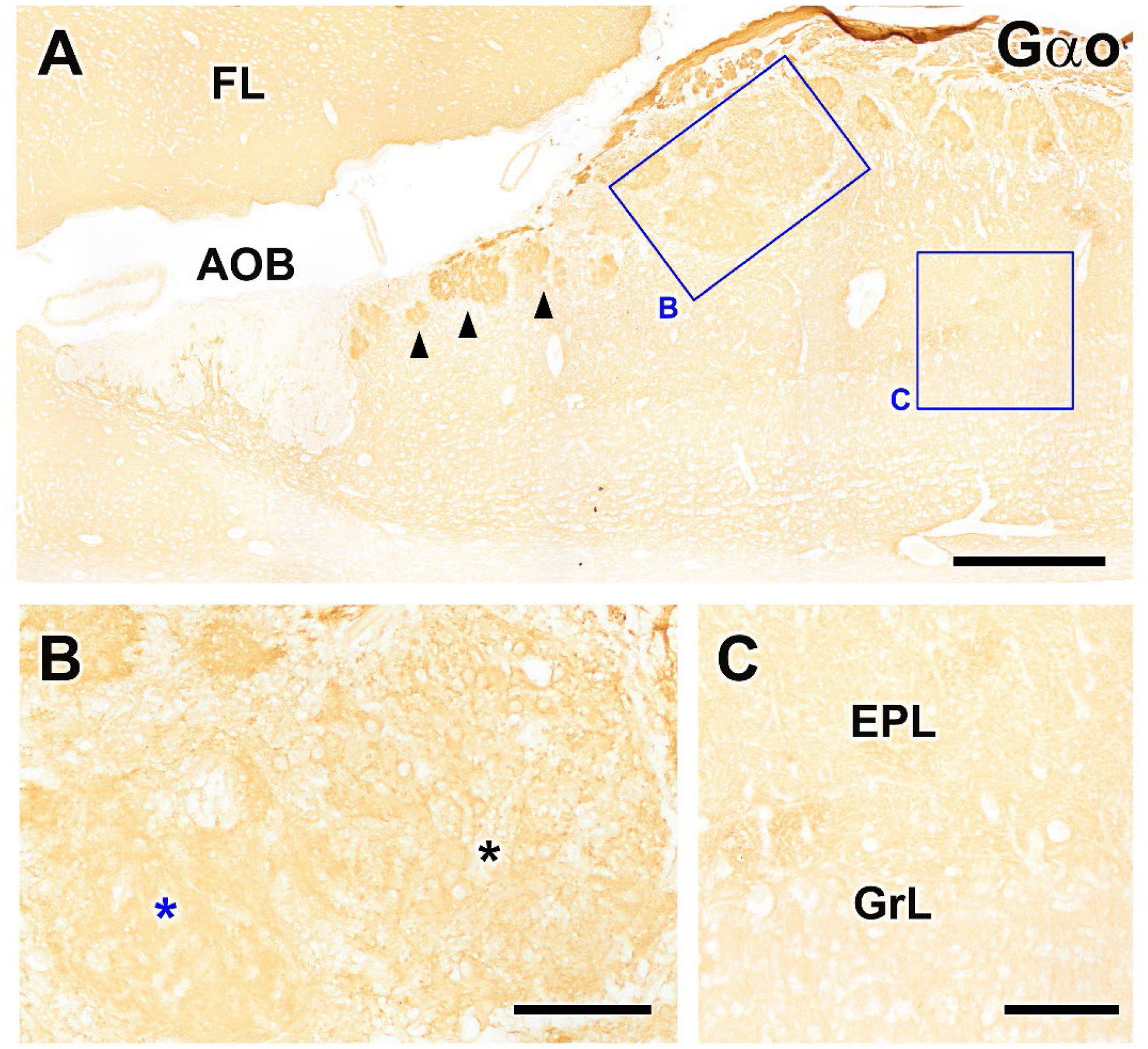
Immunohistochemical labeling of the fox olfactory limbus with anti-Gαo. **A**. The expression of the G-protein subunit Gαo produced a uniform immunopositive labeling in the frontal lobe of the telencephalon (FL) and the olfactory bulb, with only absence of immunostaining in both the nerve and glomerular layer of the AOB. However, the atypical glomeruli, close to the AOB (arrowheads), are more intensely immunolabeled than the typical glomeruli of the MOB. The MGC (box B) shows an intensity of labeling similar to the typical MOB glomeruli. **B**. Higher magnification of the box in A, showing how the MGC possess two clearly differentiated areas; one consisting of numerous neuronal somata (black asterisk) and the other devoid of them (blue asterisk). These somata are immunopositive for Gαo, which contrasts with the somata of the mitral cells of the MOB, which are immunonegative, as it is shown in the box **C**. EPL, external plexiform layer; GrL, granular layer. Scale bars: A: 500 µm. B, C: 125 µm.

Resuming the study of the OL by the double immunolabeling showed in Fig. 1, at a higher magnification in Figs. 7A and B it can be seen how the nerve and glomerular layers of the AOB are intensely labeled by the lectin, whereas the MOB glomeruli are not labeled. In the olfactory limbus some glomeruli are negative for the lectin, whereas the rest, mainly smaller, are strongly labeled with UEA (Fig. 7A). Remarkably, the Gao somata belonging to the nervous formation are embedded in a dense network of UEA-positive fibers (Fig. 7B, C). It is also noticeable how the vomeronasal nerve fibers segregate into two components: one of them UEA positive (Fig. 7B, black asterisk) and the other Gao immunopositive (Fig. 7B, white asterisk).

**Figure 7.**
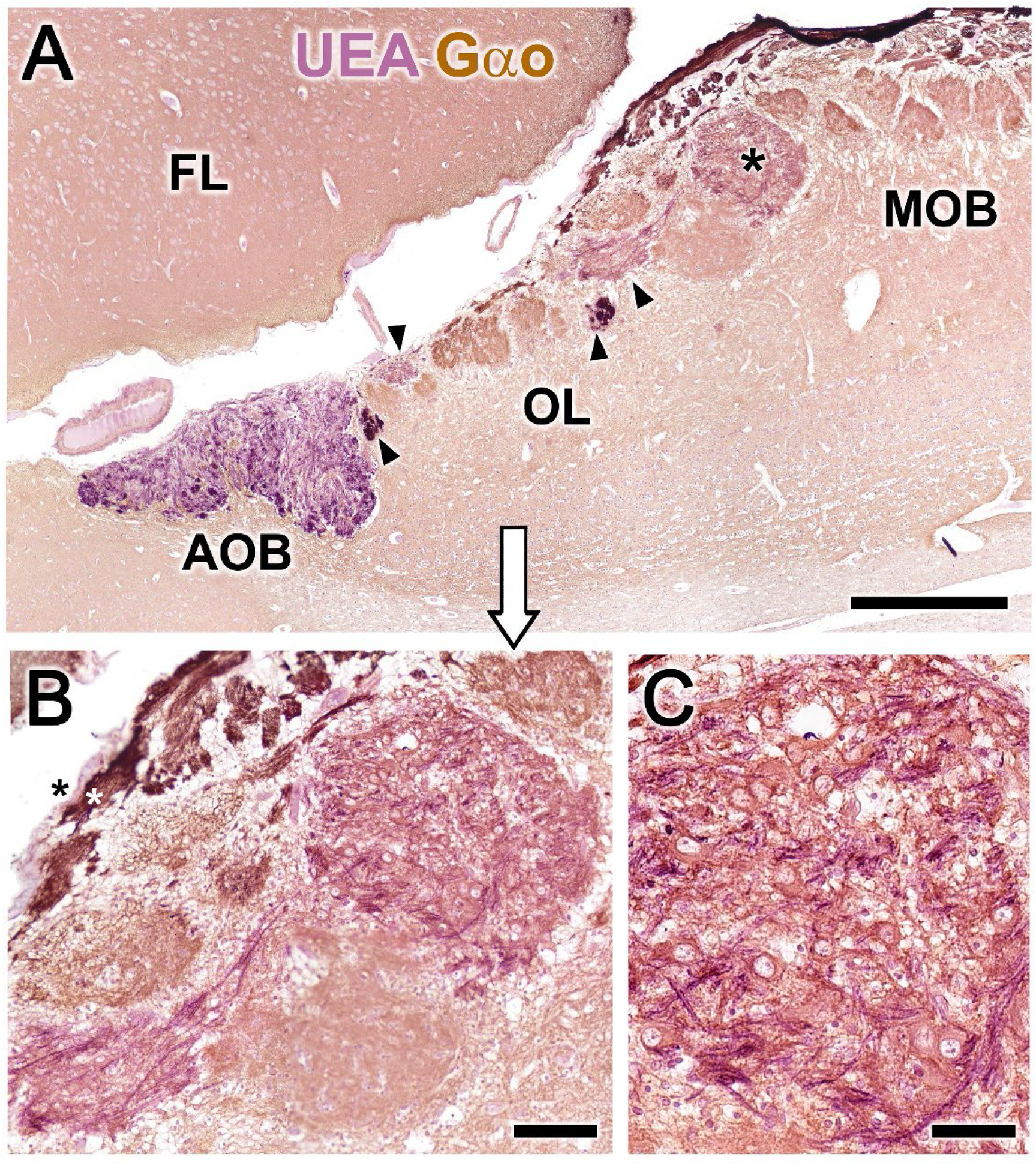
Double-immunohistochemical labeling of the fox olfactory limbus. Double immunostaining with the lectin UEA (magenta) and Gαo antibody (brown). **A**. The nerve and glomerular layers of the accessory olfactory bulb (AOB) are intensely labeled by the lectin, whereas the main olfactory bulb (MOB) is not labeled. In the olfactory limbus (OL) some glomeruli are negative for the UEA lectin, whereas the rest, mainly smaller, are strongly labeled with UEA (arrowheads). The MGC (asterisk) is also UEA positive. **B**. Higher magnification of the MGC area showing how its Gαo-positive somata are embedded in a dense network of UEA-positive fibers. It is also noticeable how the vomeronasal nerve fibers segregate into two components: one of them UEA+ (black asterisk) and the other Gαo positive (white asterisk). **C**. The somata of the MGC show an oval shape and Gαo. immunopositivity. FL: Frontal lobe of the telencephalon. Scale bars: A: 500 µm. B: 50 µm.

The anti-MAP2 antibody, which is a consistent marker for the neuronal dendritic branching of cells in all species produces a uniform immunolabeling of the glomeruli and the external plexiform layer of the MOB (Fig. 8A). In the olfactory limbus it also stains the atypical glomerular formations, whereas a patch of the MGC remains unstained (Fig. 8B, asterisk). Counterstaining of an immunolabeled section with hematoxylin (Fig. 8C) confirms that this area corresponds to the superficial area rich in neuronal bodies receiving the UEA+ innervation described in Fig. 7B.

**Figure 8.**
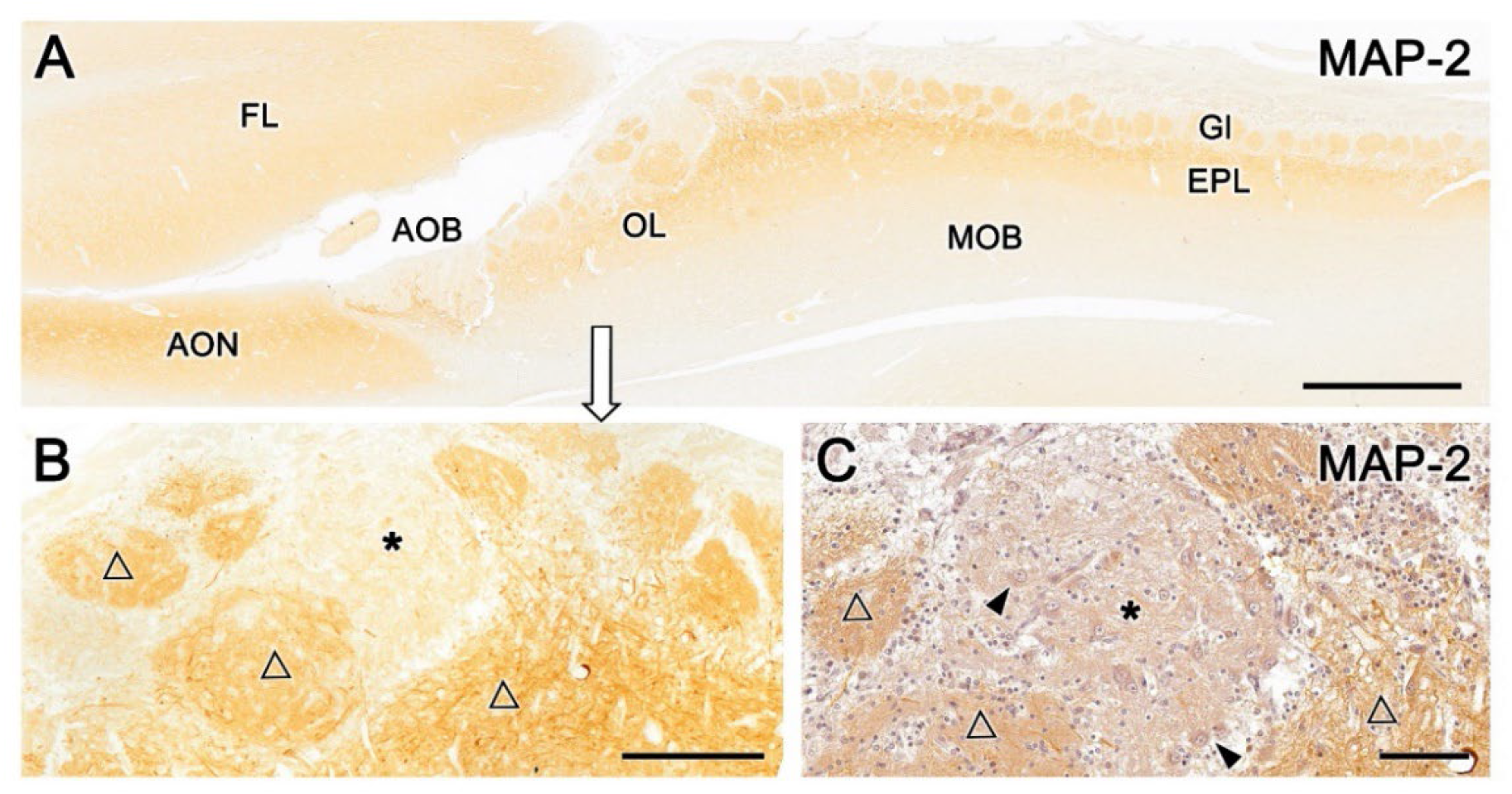
Immunohistochemical labeling of the fox olfactory limbus with anti-MAP-2. **A**. Immunostaining with anti-MAP-2 results in a strong and uniform immunolabeling of the glomeruli (Gl) and external plexiform layer (EPL) of the main olfactory bulb (MOB). **B**. A higher magnification of the rostral part of the olfactory limbus shows how the atypical glomerular formations are also stained (open triangles), but a patch of the nervous formation that remains unstained (*). **C**. An anti-MAP-2 inmunostained serial section counterstained with hematoxylin confirms that this area corresponds to the part of the MGC rich in neuronal bodies (arrowheads) that receives the UEA innervation. AOB, accessory olfactory bulb; AON, Anterior olfactory nucleus; EPL, external plexiform layer; FL, front lobe of the telencephalon; Gl, glomeruli; MOB, main olfactory bulb; OL, olfactory limbus. Scale bars: A: 1 mm. B: 250 µm. C: 100 µm.

Calcium-binding proteins are commonly used markers in the study of the olfactory bulb. Among them, we have selected three markers for the characterisation of the olfactory limbus: anti-calretinin (CR), anti-calbindin (CB) (Fig. 9) and anti-secretagogin (SG) (Fig. 10).

**Figure 9.**
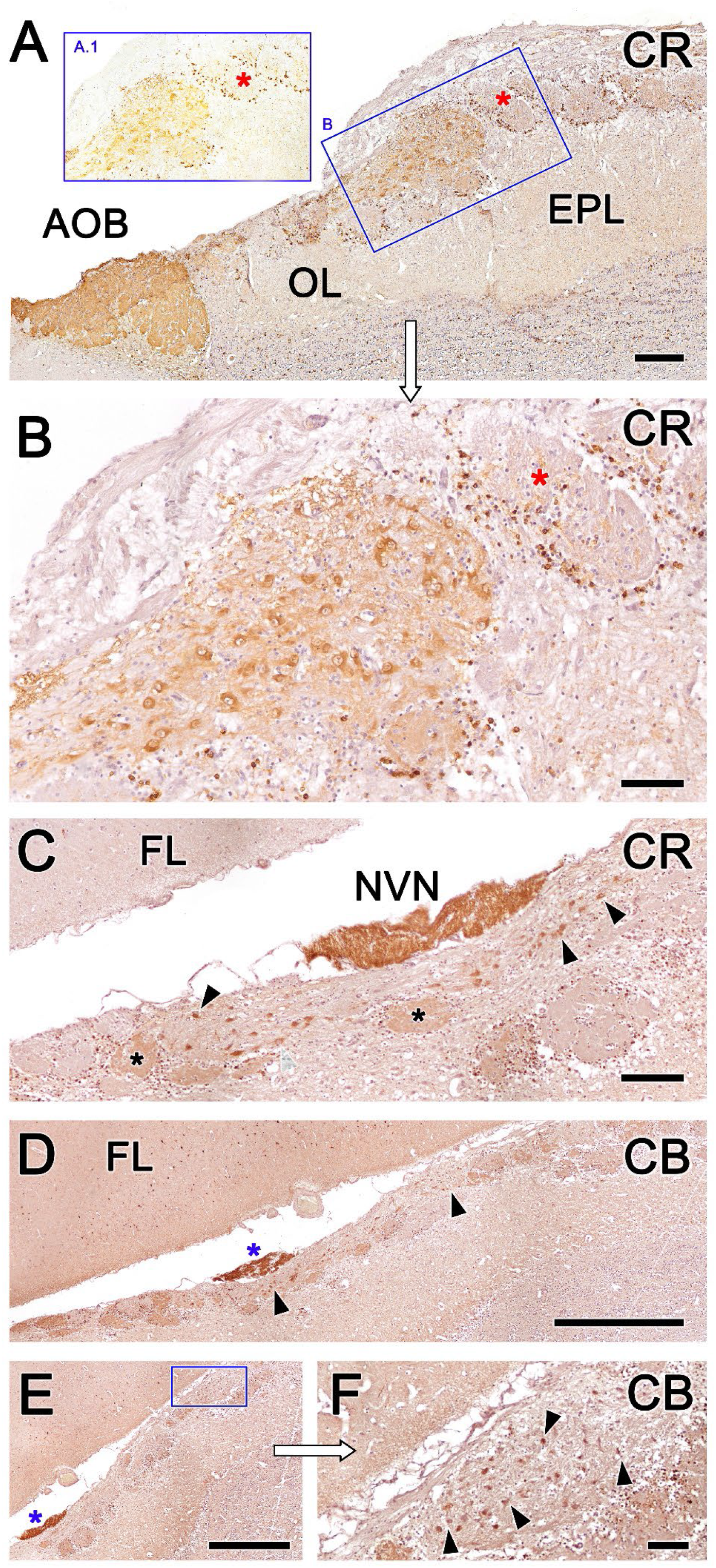
Immunolabeling with calcium-binding proteins calretinin (CR) and calbindin (CB) in the fox olfactory limbus. Hematoxylin counterstained sections. **A**. Anti-CR produces intense labeling in the MGC of the olfactory limbus (box B) as well as in the AOB. The glomeruli of the MOB do not show any immunolabeling in their neuropil (red asterisks), although their periglomerular cells are clearly labeled, what is particularly evident in non-counterstained sections (box A.1). **B**. The MGC framed in A is shown at higher magnification. The immunolabeling comprises both the neuronal somata and the neuropil (asterisk). **C**. Section performed at the level of the vomeronasal nerve (NVN), intensely immunopositive to anti-CR. The atypical nervous formation beneath the NVN is elongated and it contains scattered neuronal somata (arrowheads). A subpopulation of glomeruli is calretinin positive (asterisks). **D**. Anti-calbindin produces a similar pattern to anti-CR. The NVN is intensely immunopositive to anti-CB (blue asterisk). The MGC is delimited by arrowheads. **E**. Atypical glomeruli are variably labeled with anti-CB. **F**. Higher magnification of the inset in E. Somata belonging to the MGC are intensely stained (arrowheads). AOB, accessory olfactory bulb; OL, olfactory limbus; EPL, external plexiform layer; FL, frontal lobe of the telencephalon; NVN, vomeronasal nerve Scale bars: D, E: 1 mm. A, C: 250 µm. B, F: 100 µm.

**Figure 10.**
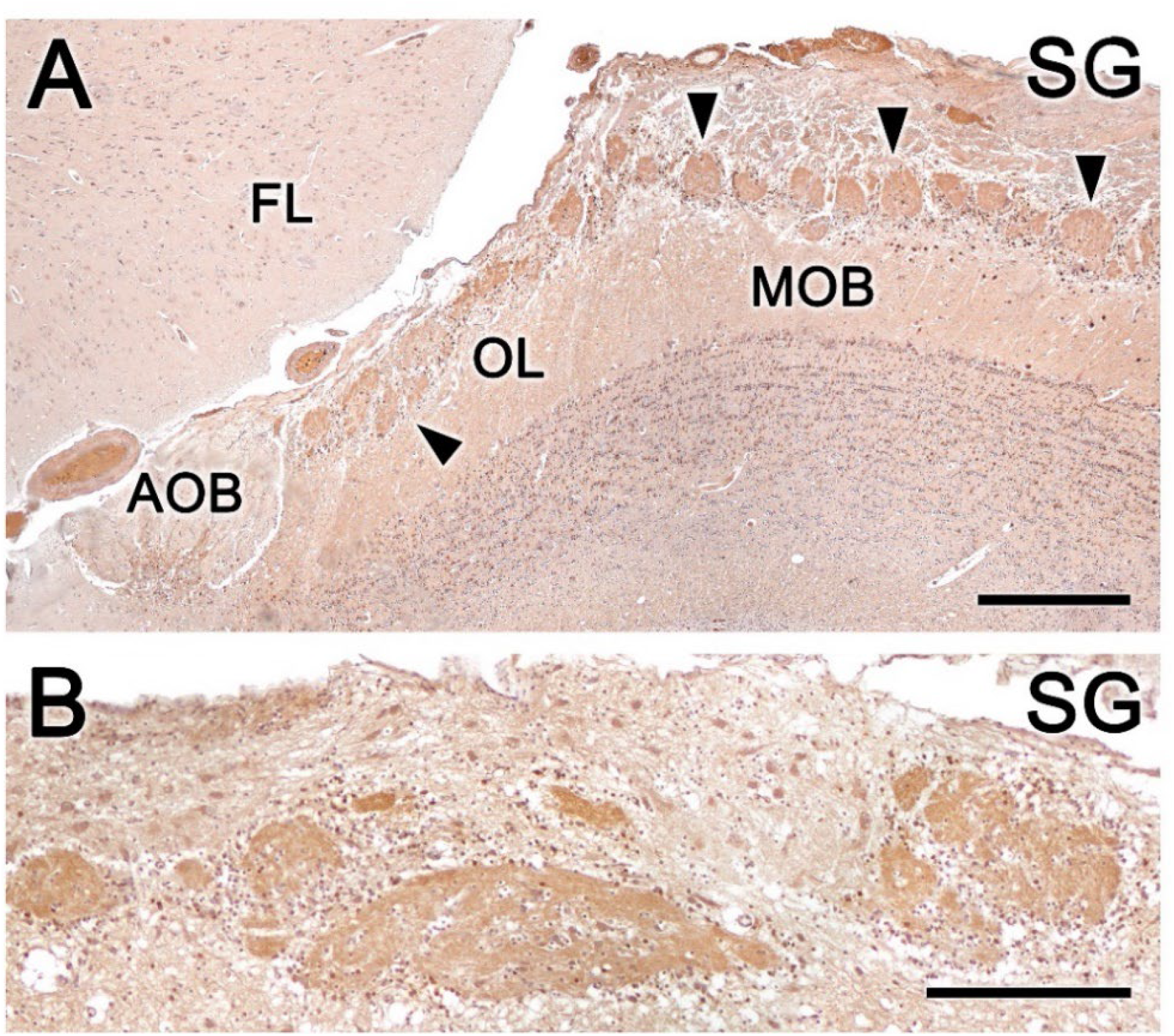
Immunolabeling with calcium-binding protein secretagogin (SG) in the fox olfactory limbus. Hematoxylin counterstained sections. **A**. Anti-SG immunolabels the whole glomeruli population in both the olfactory limbus (OL) and the main olfactory bulb (MOB) (arrowheads), but it does not stain the accessory olfactory bulb (AOB) glomerular layer. Periglomerular cells are intensely labeled. **B**. In the MGC both the neuropil and the neurons are SG-immunopositive, the latter in a variable labeling intensity. FL, frontal lobe of the telencephalon. Scale bars: A: 500 µm. B: 250 µm.

Anti-CR produces intense labeling in the MGC of the olfactory limbus as well as in the AOB (Fig. 9A). However, the glomeruli of the MOB do not show any immunolabeling in their neuropil, although their periglomerular cells are clearly labeled (insets A.1 & B in Figure 9A). In the MGC, the immunolabeling comprises both the neuronal somata and the neuropil (Fig. 9B). It is remarkable that anti-CR does not stain the mitral somata of the MOB. In sections performed at the level of the vomeronasal nerve and in which the MGC is elongated and the neuronal somata are dispersed, the immunopositive pattern described above is maintained (Fig. 9C). At this level not all atypical glomeruli are calretinin positive.

The immunolabeling pattern obtained with anti-calbindin is similar to that observed with anti-CR. Somas belonging to the MGC are intensely stained and atypical glomeruli are variably labeled, thus differentiating CR+ and CR-subpopulations of atypical glomeruli (Fig. 9D-F).

Anti-secretagogin produces a different immunolabeling pattern, immunostaining the whole glomeruli population, both in the olfactory limbus and in the main olfactory bulb. Likewise, periglomerular cells are intensely labeled (Fig. 10A). Both the neuropil and the neurons scattered in the MGC are immunopositive for SG, although in a variable degree of labeling (Fig. 10B).

## DISCUSSION

Much of the focus of investigation on the olfactory bulb has been concerned with the MOS and the VNS. Despite substantial advancements in our understanding of other subsystems, most specially the necklace region, ring of interconnected glomeruli that encircles the caudal end of the MOB and the anterior AOB, the majority of these research has been performed on rats and mice. The presence and organisation of olfactory subsystems in other mammalian groups has been very poorly studied. However, the comparative neuroanatomy studies of the MOB and AOB of laboratory rodents and other mammals such as lagomorphs (Villamayor et al., 2020), bats (Frahm and Bhatnagar, 1980), canids (Choi et al., 2010; Ortiz-Leal et al., 2022a), artiodactyls (Park et al., 2014; Kondoh et al., 2017b), and primates (Alonso et al., 1998) have found substantial differences between orders. It is therefore conceivable that at the integrative level of olfactory subsystems there are still undetected non-canonical morphological patterns which should be thoroughly addressed in order to comprehend the olfactory physiology from a solid and reliable morphological basis. This study provides neuroanatomical, immunohistochemical and lectin histochemical evidence of the presence in the olfactory limbus of a wild Canidae model species, the fox, of a complex anatomical and morphofunctional organization. This finding opens the door to additional research on this area in other mammalian species, including humans, as the olfactory limbus has thus far been poorly defined in most mammals.

### The histology of the olfactory limbus of the fox

The serial histological study of the olfactory limbus in foxes allowed us to determine in all the animals examined the presence of a complex glomerular organisation that extends along the area located between the caudal end of the MOB and the anterior extremity of the AOB. As a precedent of this atypical organisation in canids, to our knowledge, we can only cite the work of Miodonski (1968) and Nakajima et al. (1998) carried out in the olfactory bulb of the dog. The unusual organisation described by both authors in the transition zone between the AOB and the MOB does not go beyond a population of atypical glomeruli, linked for the former author to the AOB and for the latter to the processing by the MOB of information from a subset of olfactory receptors. However, in our serial study we have found in the fox a complex organisation that goes beyond the mere presence of atypical glomeruli, in terms of their size and their neurochemical and lectin-histochemical properties. Thus, the presence of neuronal aggregates comprising large somata and nascent prolongations clearly visible with Nissl staining is remarkable. These somas may be organised either in compact aggregates with a high neuronal density (Fig. 2A.1, 2B.1, 3A.1) or scattered along superficial nervous formations that extend over several millimetres that are made up of a broad neuropil formed by bundles of nerve fibres, and that we have termed as macroglomerular complex (MGC) (Fig. 2A.2, 2B.2; 3A,B,C; 4A). The presence of these neuronal clusters in such a superficial position in the olfactory bulb is a fact, to our knowledge unpublished and surprising, because even in the case of rodents, the existing descriptions of the necklace complex (Ring et al., 1997; Luo, 2008), the atypical glomerular formations (Giannetti and Le Jeune, 1996; Gómez et al., 2005) and of the olfactory limbus itself (Vargas-Barroso et al., 2017) do not contemplate the presence of superficial clusters of neuronal somata of this nature or similar.

Only a certain analogy could be established with the sub-bulbar formations described by Larriva-Sahd in the rat (2012) and Villamayor et al. (2020) in the rabbit, linked by the former with the anterior portion of the anterior olfactory nucleus and with the AOB by the latter. These formations are made up of clusters of large and polygonal neurons, but are differentiated by their location in the deepest part of the caudal part of the OB in direct association with the lateral olfactory tract to which they incorporate their axons.

### Neurochemistry of the olfactory limbus of the fox

The immunohistochemical and lectin analysis has allowed us to study in depth the neurochemical characteristics of these neurons and their associated neuropil. The most relevant fact is the immunopositivity both in the somas and in the neuropil of the atypical glomeruli against the αo subunit of the G protein, a protein widely expressed in the olfactory bulb, but absent in the somata of the mitral cells of the fox MOB. This feature establishes an important difference between the two systems and rejects the possibility that we are dealing with ectopic clusters of the MOB. No less significant is the extensive UEA+ innervation in the macroglomerular complex, observable by double immunohistochemistry (Fig. 7B). The UEA lectin is an excellent histochemical marker of the α-fucose (Kondoh et al., 2017a). In addition to mediating processes such as learning and memory, as well as neurite outgrowth and synaptic plasticity (Kalovidouris et al., 2005; Matthies et al., 1996; Murrey et al., 2009), α-fucose is an excellent marker in several mammalian species of the vomeronasal pathway, such as dogs and pigs (Salazar et al., 1994, 2000). We recently verified this fact in our study of the fox AOB (Ortiz-Leal et al., 2022b), but now, the specific study of the olfactory limbus shows us how the UEA+ nerve endings coming from the vomeronasal nerve, not only innervate the AOB but also a subpopulation of atypical glomeruli of the olfactory limbus and the Gαo+ neuronal clusters of the MGC (Fig. 7B).

Anti-MAP-2 is a useful marker for mitral/principal cells dendritic trees (Dehmelt and Halpain, 2005) and it is absent from axons (Bernhardt and Matus, 1984). Its use in the olfactory limbus is therefore a good tool to characterize the shape of its glomeruli. We have found that atypical glomeruli stain more intensely than MOB glomeruli and show a greater morphological diversity (Fig. 6A). This has been previously demonstrated by morphometry in the mouse glomerular necklace complex (Walz et al., 2007). Another differential aspect between the olfactory limbus and the MOB is that the MGC somata stain more intensely than those of the MOB mitral cells (Fig. 6B).

The investigation with calcium-binding proteins, markers widely used in the study of OB (Crespo et al., 1997; Defteralı et al., 2021), also allows to establish neurochemical differences between the limbus and the main olfactory bulb. Calretinin and calbindin intensely mark the atypical glomeruli and neuronal aggregates of the olfactory limbus, in both cases at both the neuronal somata and their neuropil. However, in the case of MOB glomeruli, there is hardly any labeling in the neuropil, although there is labeling in the periglomerular cells. The presence of abundant periglomerular CB and CR immunopositive cells in the atypical glomeruli is similar to that observed in the olfactory bulb of the rat (Crespo et al., 1997). It is striking that the principal neurons of the olfactory limbus, unlike the mitral cells of the fox main olfactory bulb, show positive immunolabeling against both anti-CB and anti-CR. Secretagogin is a more recently discovered calcium-binding protein (Wagner et al., 2000) and therefore less studied in the olfactory bulb, where it is widely expressed throughout the OB layers (Kosaka and Kosaka, 2013; Pérez-Revuelta et al., 2020). In the case of the fox, this broad pattern of immunolabeling is conserved in both the olfactory limbus and the main olfactory bulb, but not in the accessory olfactory bulb.

### Functional correlates for the fox olfactory limbus

The innervation of the olfactory limbus neuronal clusters by UEA+ fibres establish a clear morphofunctional link between the olfactory limb and the vomeronasal system, as UEA lectin is a highly specific marker in the vomeronasal system of the fox: vomeronasal neuroepithelium, vomeronasal nerves and the nerve and glomerular layers of the AOB. To this must be added the Gαo character of the neuronal clusters of the OL, both at the level of their somas and the neuropil. This fact constitutes an important functional differentiation with respect to the main cells of the MOB, in addition to the different neurochemical patterns identified with antibodies against MAP-2, CB and CR between the main cells of the OL and the mitral cells of the MOB.

These morphological and neurochemical observations open the possibility of linking functionally the olfactory limbus with the extensive Gαo immunoreactivity found in the neuroepithelium of the fox vomeronasal organ (Ortiz-Leal et al., 2020). Subsequent neurochemical characterisation of the fox AOB (Ortiz-Leal et al., 2022b) failed to detect the presence of Gαo immunoreactivity in the nerve layer of the AOB leading to the conclusion that sensory information detected by the vomeronasal receptors associated with the fox Gαo should project to a different bulbar territory. Therefore, the functional linkage of OL with UEA+ fibres from the VNO, the high OL-associated Gαo immunoreactivity, as well as the presence of Gαo+ axons in direct topographical relation with OL and the vomeronasal nerve (Fig. 7B) make plausible the hypothesis that the olfactory limbus is at least partially involved in the processing of sensory information from the Gαo neuroepithelial cells of the VNO.

The connection of the fox olfactory limbus with the vomeronasal system and its strategic position between the main and accessory olfactory bulbs suggests that the fox’s OL is involved in the processing of specific stimuli that signal relevant intraspecifical socio-sexual cues in line with what has been suggested by other authors in laboratory rodents (Weruaga et al., 2001; Leinders-Zufall et al., 2007; Larriva-Sahd, 2012; Vargas-Barroso et al., 2017).

### Canids domestication and olfactory limb morphology

The high degree of development and structural and neurochemical complexity found in the olfactory limbus of the fox raises the question of why in the dog, as the most studied canid, no similar structure or organisation has been found in this region. Only the aforementioned studies by Miodonski (1968) and Nakajima (1998) point to the presence of atypical glomerular organisations in the dog, but in neither case are they comparable to those observed in the fox.

The fox is a particularly good model for researching the impacts of domestication in Canidae due to a long-term experiment aimed to replicate early mammalian domestication in this species (Belyaev et al., 1985; Wang et al., 2018). Fox selection for tameability led to changes similar to those observed in the domestic dog’s behavior, physiology, and genetic diversity as a result of domestication (Kukekova et al., 2018; Trut et al., 2009).

Our previous studies of the fox VNS opened the possibility that the striking structural differences observed between the dog and the fox AOB may partially be the result of the domestication process (Ortiz-Leal et al., 2020; 2022b). The current theory that domestication of dogs has led to an involution of olfactory development linked to the detection of pheromones and other semiochemicals (Jezierski et al., 2016) is supported by the anatomical differences found in the OL of the fox compared to the data available in the dog OB.

